# *Ingres*: from single-cell RNA-seq data to single-cell probabilistic Boolean networks

**DOI:** 10.1101/2022.09.04.506528

**Authors:** Pedro Victori, Francesca M. Buffa

## Abstract

**Motivation:** The current explosion of ’omics data has provided scientists with an unique opportunity to elucidate the inner workings of biological processes that remained opaque. For this, computational models are essential. Gene regulatory networks (GRN) have long been used as a way to integrate heterogeneous data into a discrete model, and are very useful to generate actionable hypotheses on the mechanisms governing these biological processes. Boolean networks are particularly popular for this kind of discrete models. When working with single-cell RNA-seq datasets a main focus of analysis is the differential expression between subpopulations of cells. Boolean networks are limited in this task, since they cannot easily represent different levels of expression, confined as they are to binary states. We set out to develop an algorithm that can fit Boolean networks with this kind of data, maintaining both the heterogeneity of the data and the simplicity and computational efficiency of these type of networks.

**Results:** Here we present *Ingres* (**I**nferring Probabilistic Boolean **N**etworks of **G**ene **R**egulation Using Protein Activity **E**nrichment **S**cores) an open-source tool that uses single-cell sequencing data and prior knowledge GRNs to produce a probabilistic Boolean network (PBN) per each cell and/or cluster of cells in the dataset. *Ingres* allows to better capture the differences between cell phenotypes, using a continuous measure of protein activity while still confined to the simplicity of a GRN. We believe *Ingres* will be useful to better understand the heterogeneous makeup of cell populations, to gain insight into the specific circuits that drive certain phenotypes, and to use expression and other omics to infer computational cellular models in bulk or single-cell data.

**Availability and implementation:** *Ingres* has been implemented as an R package, and it is publicly available at https://github.com/CBigOxf/ingres. It is currently being submitted to the public repository CRAN too. works seamlessly with existing software for single-cell RNA-seq analysis, and for network analysis, modelling and visualization.

**Supplementary information:** Supplementary data are available online. Software documentation and an explanatory vignette are available on GitHub and as part of the R package.

## Introduction

Biological processes are characterized by complex molecular interactions that have been modelled using a number of different frameworks. Boolean gene regulatory networks (BGRN) are a powerful tool for this purpose due to their simplicity and computational properties (Schwab et al. 2020). These qualitative models allow to distill the highly complex and stochastic nature of such processes into a model that is simple to understand and interpret. In these networks, all components —be it genes, proteins, regulatory interactions or a mixture of those— are modelled as nodes governed by binary functions, so that at any given time each node has one of two possible states (abstracted as on/off, or active/inactive).

BGRN are directed graphs, where the relationships between these nodes are expressed as logical Boolean functions. Thus, the state of each node depends on the state of other nodes. These networks can also have input nodes, which state is externally set: they do not depend on the state of other nodes within the network. They represent the agents foreign to the specific process that it is being modelled, such as a receptor protein becoming activated, an immune response being triggered, or a nutrient being absorbed by the cell. Conversely, there can be nodes with no outgoing links to other nodes, and they will be considered the output nodes, or nodes that represent the overall phenotype of the network.

One limitation of BGRNs is that they lack the ability to express the stochastic nature of biology. In such a network, be it synchronous or asynchronous (Garg et al. 2008), it is assumed that all interactions take the same time and are equally likely to happen. Probabilistic Boolean networks (PBN) can address this limitation (Shmulevich et al. 2002). In these networks, each node behaviour is described by more than one Boolean function. Whenever a node state has to be updated, one of those functions is chosen according to a given probability. The probability for all functions of a node sums up to 1. In this framework, the outcome of the network —its phenotype— is probabilistically, rather than deterministically, dependent on its previous state. This behaviour more closely resembles stochastic biological processes, while maintaining the underlying simplicity of Boolean networks. This is one of the reasons we developed *Ingres*, a method to produce PBNs using gene regulatory networks and transcriptomics data.

Another limitation of Boolean network modelling, which we also address with *Ingres*, is the simulation of continuous biological variables. For example, different levels of gene expression between cell sub-populations, or in cells exposed to different conditions, are not easily modelled in Boolean networks. Traditionally, over- and under-expression are represented by imposing a constraint to the node, fixing it to a single state throughout the simulation. The Boolean function for those nodes will be ignored and its state will always be 1 or 0, regardless of the state of connected nodes. Of course, this is the trade-off of discretisation, which is normally thought to be offset by the advantages it provides: an easier and faster computational analysis, homogenisation of different data sets and reduction of noise (Gallo et al. 2016).

On the other hand, quantitative and hybrid models have been proposed to represent biological networks, and they allow the simulation of continuous levels of expression, but they are only feasible for very small networks, since they require an extensive knowledge and experimental data to be developed (Saadatpour and Albert 2016).

*Ingres* provides another solution to this problem by representing different levels of activation/expression while still working with Boolean functions. Specifically, we use a previously developed algorithm, VIPER (Alvarez et al. 2016), to infer protein activity starting from a gene expression matrix and a list of regulons —defined as the list of transcriptional targets of each protein. This computes a matrix of normalised enrichment scores (NES) for each protein. The expression of these regulons represent an optimal indirect method to measure the activity of a specific gene/protein (our nodes in the network). This constitutes a novel way to fit the PBN using single-cell RNAseq data.

BGRNs are increasingly used also in combination within more complex, multi-scale models, such as our recently published framework microC (Voukantsis et al. 2019). These are models where each cell (e.g. modeled as an independent entity using Agent Based Modelling) is simulated as having their own gene network whose output governs the behaviour of the cell. External signals, nutrients or interactions with other cells can also be received by the cell, and act as inputs of its inner network, therefore simulating tumour dynamics at several levels: the microenvironment level, the cellular level with its cell-cell interactions and the intracellular level. These multi-scale models are a novel and exciting development in the field, and others have been proposed since microC (Letort et al. 2018). They allow to start from basic assumptions, and generate simulations to explore the emerging behaviour and evolution of heterogeneous populations of interacting cells. For this kind of models, finding a way to represent the differences between cells and between subpopulations is crucial, especially for cases where cellular heterogeneity and competition have a significant role in driving the biology of the system (Dagogo-Jack and Shaw 2018). Furthermore, the increasing availability of high-throughput screens is a tremendous training opportunity to initialize the cells and to test the hypothesis generated by the model. However, currently there is not a methodology to easily do this. *Ingres* allows not only to build BGRNs specific to an RNA-seq dataset, but also to initialise each single simulated cell’s BGRN using single-cell RNA-seq data. This opens the opportunity to model heterogeneous cell populations, and generating simulations and results that better represent specific biological and clinical contexts.

To conclude, we hope that *Ingres* will help bridge the gap between highly dimensional transcriptomic data and discrete computational models, providing a useful tool for generating mechanistic explanations and identifying actionable targets for therapy, tasks for which these models are particularly suited (Victori and Buffa 2018). To enable use from the wider community, we have included functionality so that works seamlessly with related software and algorithms such as *Seurat* (for single-cell RNA-seq analysis) or *BoolNet* and *tidygraph* (for network analysis, modelling and visualization).

## Algorithm, implementation and results

First, we will describe the steps in the *Ingres* workflow, then we will provide some examples and results. We have implemented *Ingres* as a R package, and the documentation attached to it contains specific information about the functions that are used to run each of the steps described below.

### 1. Inferring protein activity using VIPER

RNA levels are often not a good proxy for protein activity. For example, a gene function could be modified by altered translation or protein degradation, without affecting RNA levels. In other words, RNA levels do not paint the full picture of biological mechanisms, and both RNA and protein data need to be considered. However, this is often ignored in network studies which tend to be based either on protein or RNA data.

Here, we address this point by employing Virtual Inference of Protein-activity by Enriched Regulon analysis (VIPER), which wraps an algorithm called *aREA*, standing for analytic rank-based enrichment analysis. VIPER infers protein activity by analysing the gene expression of the protein’s regulon (the list of transcriptional targets of each protein), which is tissue-dependant (Alvarez et al. 2016). Instead of simply using the gene expression values to reconstruct or initialize out network, we propose to use these scores to infer protein activity. This is crucial to retain some of the information that is necessarily lost not only with the discretisation of the data, but also by retaining only the expression scores for the few dozen genes contained in a GRN. Since VIPER takes into account all genes that are targets of the proteins of interest —the ones contained in the GRN— using NES instead of gene normalised expression preserves more information, and therefore makes the resultant model more representative of the biological reality that is being studied.

VIPER probabilistically integrates the target mode of regulation (activation or repression), statistical confidence in regulator-target interactions and target overlap between different regulators (Alvarez et al. 2016). VIPER is also very robust and resilient to signature noise and to incomplete regulons (Alvarez et al. 2016). In *Ingres* we use VIPER to infer protein activity status from the provided gene expression matrix, using the regulon supplied by the user. This regulon can be inferred using methods such as the ARACNe algorithm (Basso et al. 2005; Lachmann et al. 2016), or can be extracted from repositories such as Dorothea (Garcia-Alonso et al. 2019), or can be the result of specific experiments.

If several regulons are provided, *Ingres* runs the metaVIPER algorithm (Ding et al. 2018) instead. This algorithm is designed to integrate multiple interactomes, not necessarily tissue-matched, as VIPER needed to work accurately. This is most valuable in single-cell RNA-seq data, due to its heterogeneity, inherent noisiness and low sequencing depth (Ding et al. 2018). Indeed, metaVIPER has been shown to reduce bias and batch effect, generating reproducible protein-activity signatures (Ding et al. 2018). Therefore, we recommend this method when using *Ingres*. Collections of tumourregulons are available at Bioconductor (Giorgi 2017).

### 2. Producing Probabilistic Boolean Networks through the Normalised Enrichment Scores

Boolean networks contain a set of variables *X* = {*x*_1_, *x*_2_, …, *x*_*n*_}, *x*_*i*_ ∈ 𝔹, called nodes. Therefore, the state of such a network at time *t* if defined by the vector 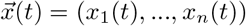. For every variable *x*_*i*_ there is a corresponding function *g* such as

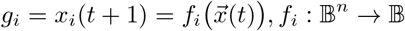

*Ingres* takes 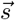, the vector containing all the NES generated by VIPER for a given cell or cluster of cells, and an user defined range such as

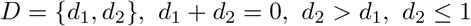

to which the scores are to be rescaled, and computes the rescaled scores (*r*) with mid-point 0 as

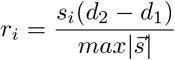

for every element of 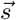, where 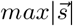 is the maximum absolute value through the NES vector. Then, in the new PBN that *Ingres* produces, to every node *x*_*i*_ where r_*i*_ ≠ 0, there corresponds a set *F*_*i*_ = {*g*_*i*_, *o*_*i*_} where *g*_*i*_ is the node function in the original Boolean network and

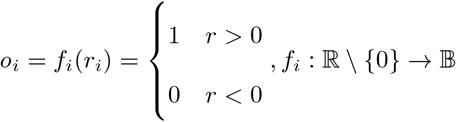

As such, this function always returns just 1 —activation— or 0 —deactivation—. For nodes *x*_*i*_ where *r*_*i*_ = 0, *F*_*i*_ = {*g*_*i*_}. For every set *F*_*i*_ there is a corresponding set of probabilities 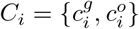, where

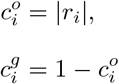

with 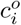 and 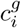 being the probabilities for the functions *o*_*i*_ and *g*_*i*_, respectively. For *F*_*i*_ with only one function *g*_*i*_ (when *r*_*i*_ = 0), 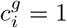. Those probabilities will be used to determine which function should be use to perform every transition *x*_*i*_(*t*) ⟼ *x*_*i*_(*t* + 1).

If the PBN is to be computed for a given cluster (a subpopulation of cells), the median NES among all cells belonging to that cluster is calculated for each gene, and then the steps above are executed taking the set of all cluster scores computed this way. This represents a useful compromise from the (large, continuous and noisy) raw expression data, and the standard rough discretisation approach of representing over-expression by fixing nodes to 1 and vice versa.

The nodes in the PBNs produced by *Ingres* will not be fixed into a single state —except for the top and bottom genes in each cell, which will have a fixed function probability of 1—, they will still be subject to their original Boolean function, but they will have a bias towards inactivation or activation directly proportional to their protein activity enrichment score (Figure 1). This bias will of course work independently from the Boolean function: at any update step where the fixed function is chosen, the incoming edges of the node will be ignored and the state will be determined by the fixed function. This simulates the action of all the factors that contribute to driving the phenotype but are not present in the GRN: other genes and proteins, external signals, environmental factors, etc., without a significant increase in complexity other than what naturally comes with switching to a probabilistic network. The user, through the rescaling range parameter (which defaults to [-1, 1], the maximum), can decide how strong this bias will be in the generated network.

**Figure 1.**
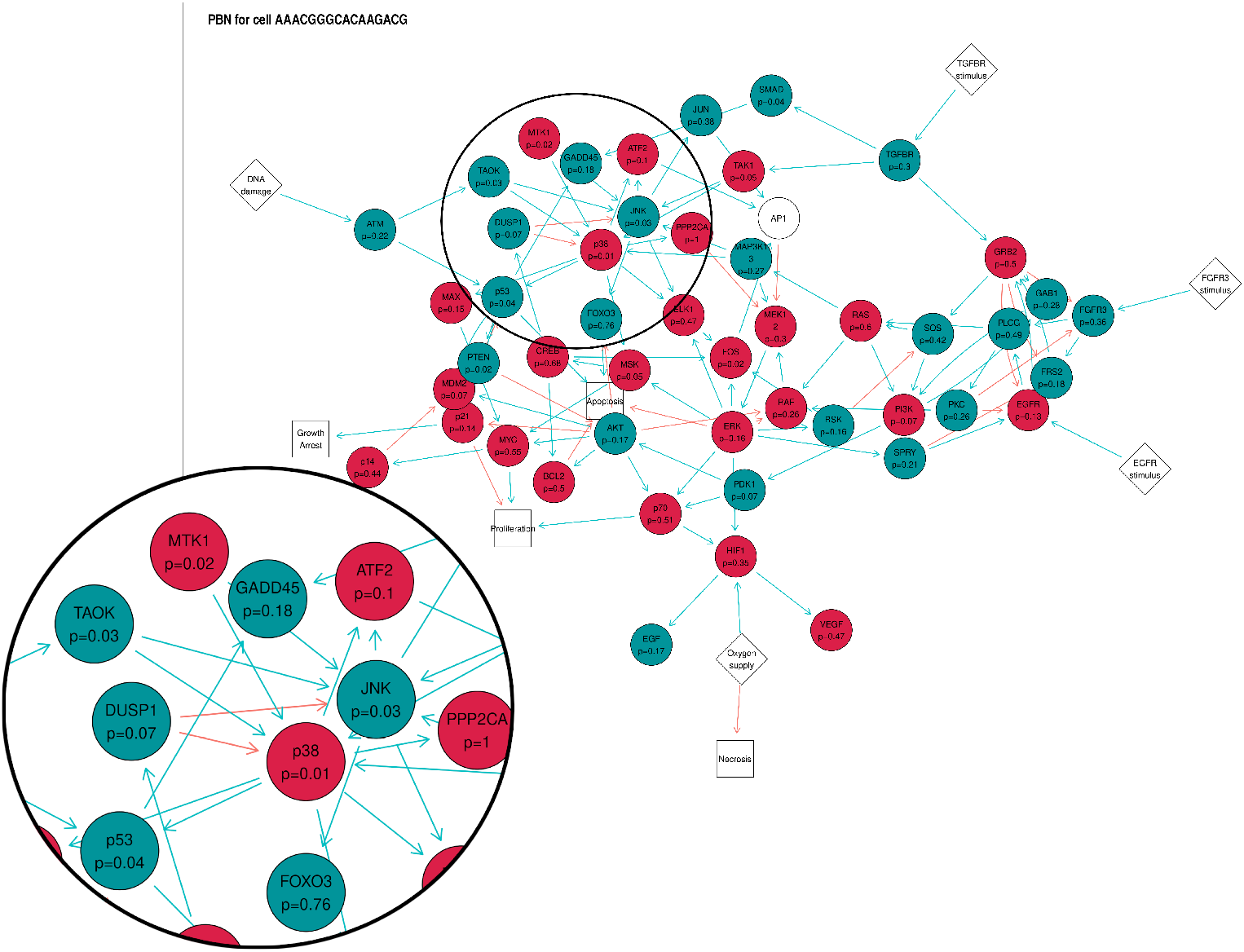
A network generated by *Ingres*. A network plot showing the probabilities for the fixed function of each node. The green nodes indicate that their fixed function is 1 (activation) and for the red nodes it is 0 (deactivation).

### 3. Visualization and analysis of the networks generated by *Ingres*

Now, the user can explore the results by using *Ingres* functions to plot the network for any cell or cluster. The results can be plotted as heatmaps too. Additionally, an interactive app can be run, where the user can see all cells in an tSNE plot — by providing a Seurat object (Stuart et al. 2019) — and click on any of them to be shown its corresponding PBN (Figure 2).

**Figure 2.**
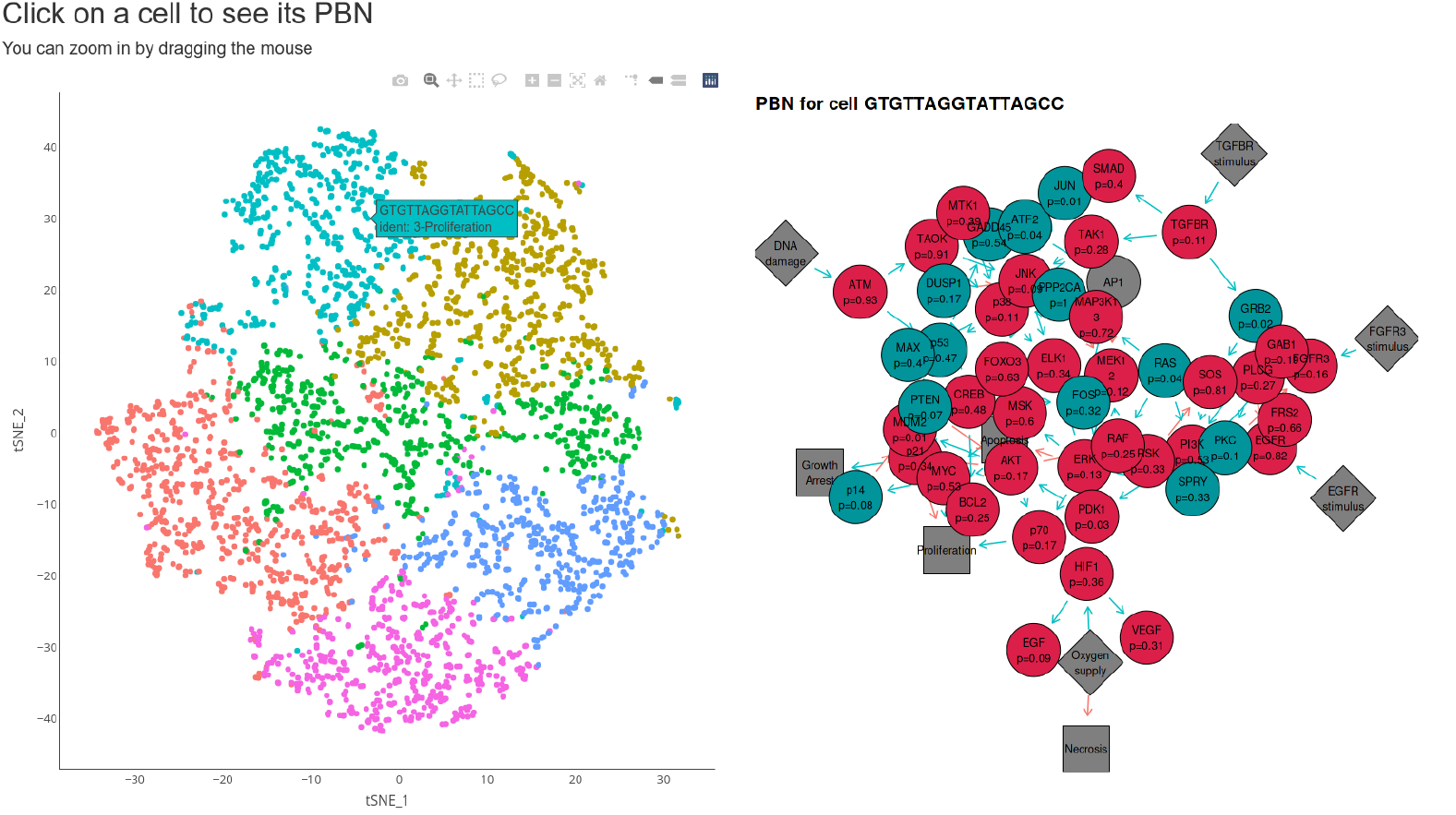
Interactive app for visual exploration of the results. Screenshot of the app that allows the user to click on any cell on the tSNE cell and shows the computed PBN of the selected cell.

Finally, *Ingres* provides several wrapper functions for relevant parts of *BoolNet* (Müssel et al. 2010), which can be used to perform analyses on any PBN produced by *Ingres*, such as computing its attractors (Hopfensitz et al. 2013). This can be useful to identify predicted phenotypes of the modelled network. *Ingres* exports its networks in *tidygraph* (Petersen 2020) and *BoolNet* format, so they can be further interrogated using other software, such as *GinSim* (Chaouiya et al. 2012).

### 4. Testing *Ingres* using a single-cell RNA-seq atlas of breast cancer cell lines

In order to provide a proof of concept and a functional validation of *Ingres*, we exploited a recently published single-cell RNAseq atlas of breast cancer cell lines (Gambardella et al. 2022). We employed a published growth factor and cell cycle signalling network (Remy et al. 2015) to infer our networks. First, we removed redundant nodes and reduced it to 30 nodes. We then processed the RNAseq data from the published single-cell atlas, obtaining data for 35000 single-cells (Gambardella et al. 2022) in 31 cell lines, which were used as the input for *Ingres*. For every cell line, we clustered cells based on their cell-cycle score, as described by the Seurat authors (Seurat 2022), in three clusters, G1, S and G2M. Finally, we performed the whole *Ingres* pipeline on each dataset, obtaining NES scores and PBNs for each single-cell (**Supplementary materials**).

We then explored the differences between the PBNs in the three clusters. We computed 1000 state transitions for each network to see what the fate activation trend is for each of them (Figures 3 and S1). As expected, as the cell cycle progresses, the networks corresponding to its phases show a increasing proportion of proliferation, and in the S and G2M networks, it is higher than apoptosis. This result is very robust across the different breast cancer cell lines in the input data. Therefore, as long as the input network is chosen based on the phenotype being studied, the PBN generated by *Ingres* preserves the expected behaviour of each of the cell clusters. The same simulation using a dataset of bladder cancer samples (Wang et al. 2020) (available as GSE130001) as the input for *Ingres* was performed with similar results (Figures S2 and S3).

**Figure 3.**
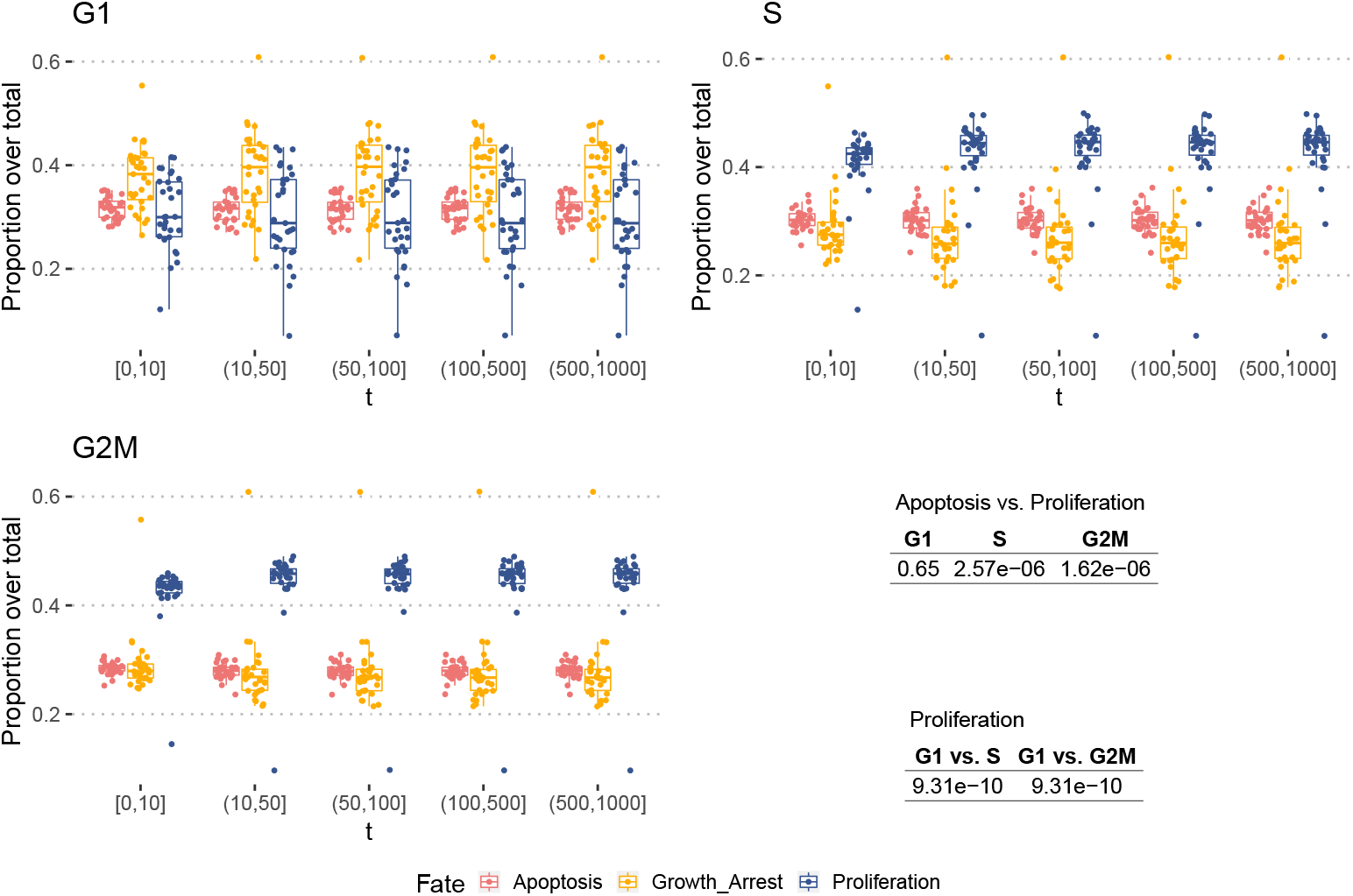
State transition.

## Discussion

In this article, we have described a new tool, *Ingres*, that can infer gene regulatory Boolean networks from single-cell RNA-seq data, producing probabilistic Boolean networks that are specific to each cell. *Ingres* allow us to infer the probability of each node being active and to compute each gene’s normalised enrichment score. In doing this, we take into account each network gene’s whole regulon instead of just the gene’s expression level, allowing to account for the global RNA-seq information into the final PBN, rather than just the expression of the single gene node. Using PBNs this way allows to model the difference of gene expression levels between cells or clusters, therefore allowing researchers to simulate scenarios that are more complex than total silencing or over-expression of genes.

We believe this approach is a good compromise between the simplicity and flexibility of Boolean networks and the wealth of information that single-cell sequencing provides. *Ingres* facilitates fitting models with cell-specific expression information without the need of inferring a new network for each cell or cluster, thus making the result more comparable from cell to cell, and scalable for application to cell-based simulations involving hundreds of cells. Indeed, the resultant networks still have discrete states and outputs, which makes them perfect candidates to use in combination with agent-based modelling of cell and tissue dynamics. Finally, while we developed this tool to aid our research on multi-scale biological modelling, we expect it will prove useful for the biology community at large to aid in the interpretation of the large quantity of high-throughput data which are being generated.

## Acknowledgements

The authors would like to thank Dr. Marta Pérez Alcántara and Professor Tim Maughan for their helpful comments.

## Author Contribution

PV and FMB conceived the idea. PV implemented the software with help from FMB. FMB supervised the work. FMB and PV wrote the manuscript.

## Supplementary data

### ingres 2022 supplementary data.zip

This folder, available for download in Zenodo (https://doi.org/10.5281/zenodo.7025732), contains the data generated by *Ingres* as described in *4. Testing Ingres using a single-cell RNA-seq atlas of breast cancer cell lines*. It consists of one “remy network 2022-05-23.rds” R data file for each of the 31 cell lines and 3 cell cycle stage. Each of them stores the corresponding network produced by *Ingres* as a BoolNet PBN. The folder also contains one “ingres-viper remy 2022-05-20.rds” R data file for each of the 31 cell lines. These files store *Ingres* R objects loaded with the corresponding cell line single-cell data and the subsequent VIPER analysis results.

## Supplementary Figures

**Figure S1.**
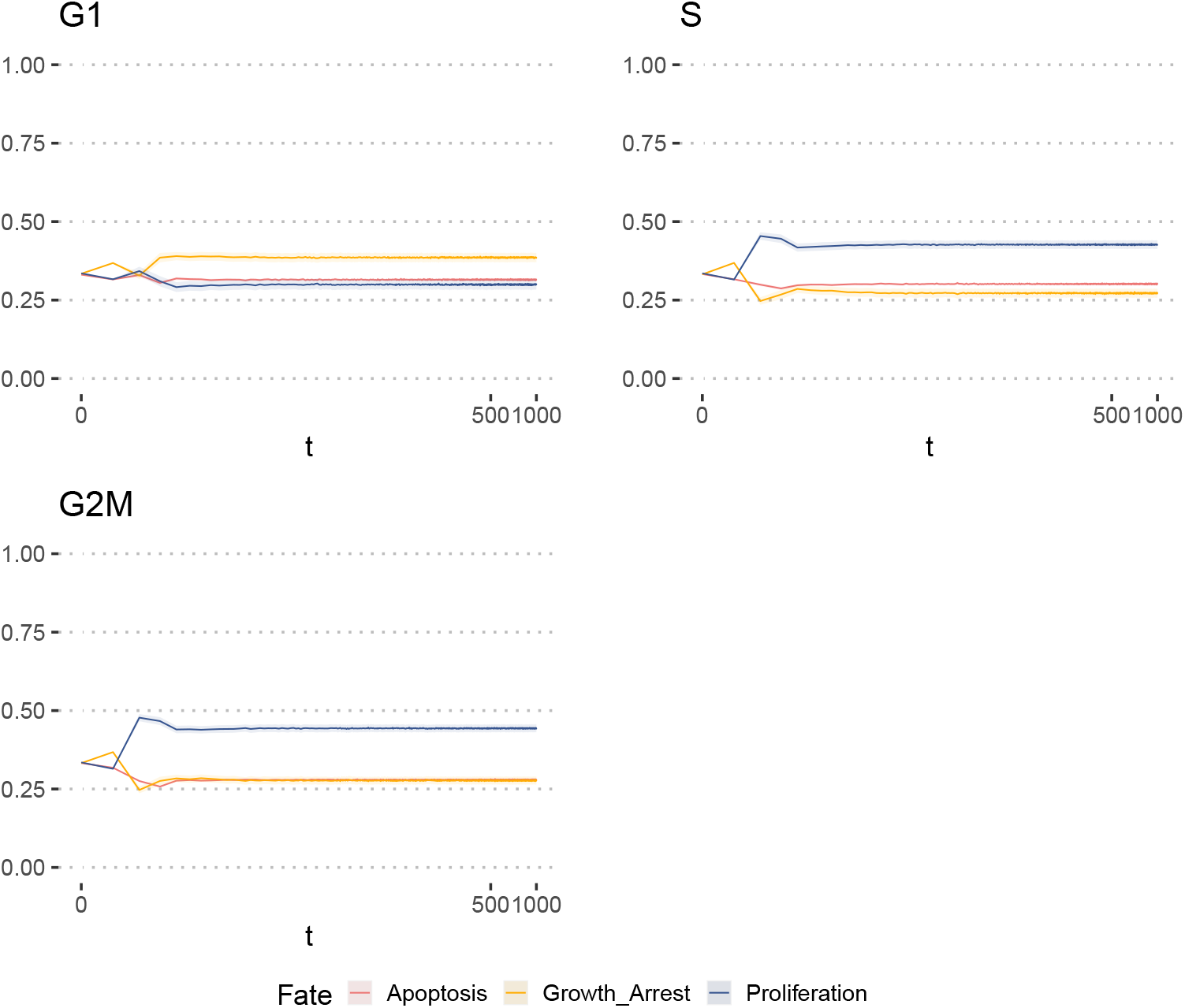
State transition computation for the networks produced by *Ingres*. The data used for this figure was produced as described in Fig. 3. The figure shows the proportion of each fate over the total at each state transition (shown with a pseudo-logarithmic scale), averaged over the 32 cell line. The plot shows the standard error of the mean too.

**Figure S2.**
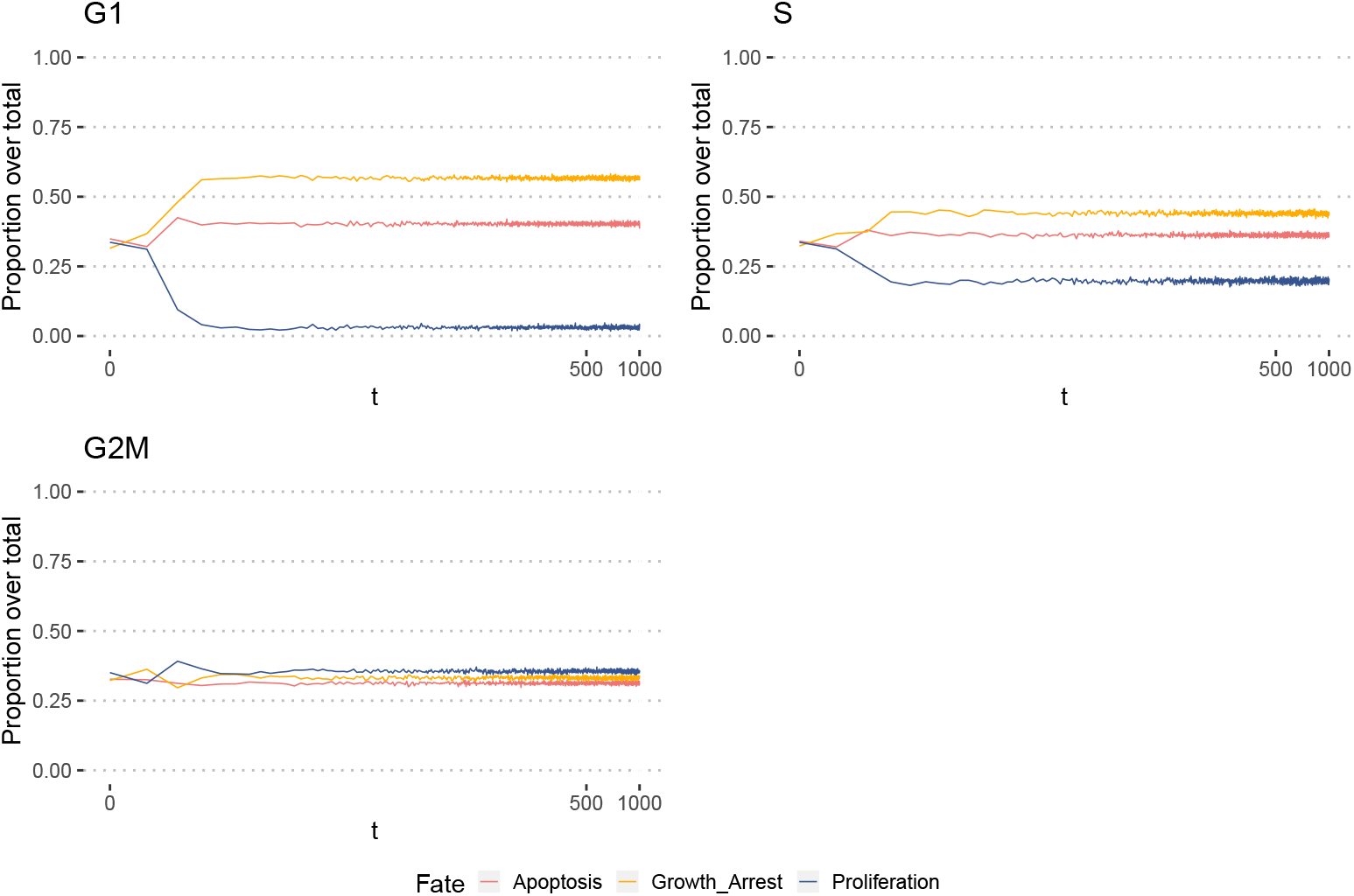
State transition computation for the networks produced by *Ingres* using a bladder cancer dataset. The procedure for this simulation was the same as described for Figure 3. The figure was produced as described for Figure S1.

**Figure S3.**
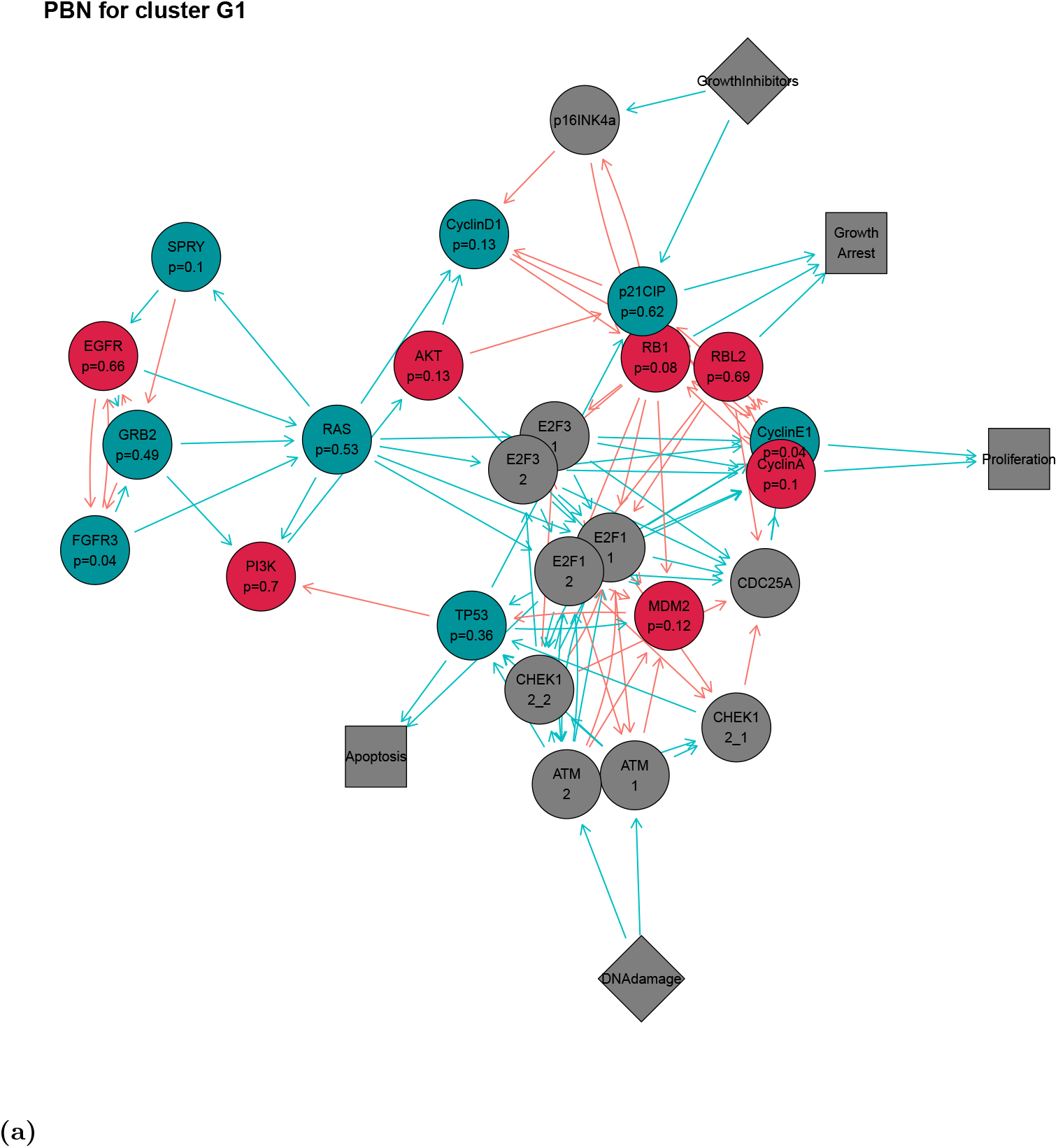
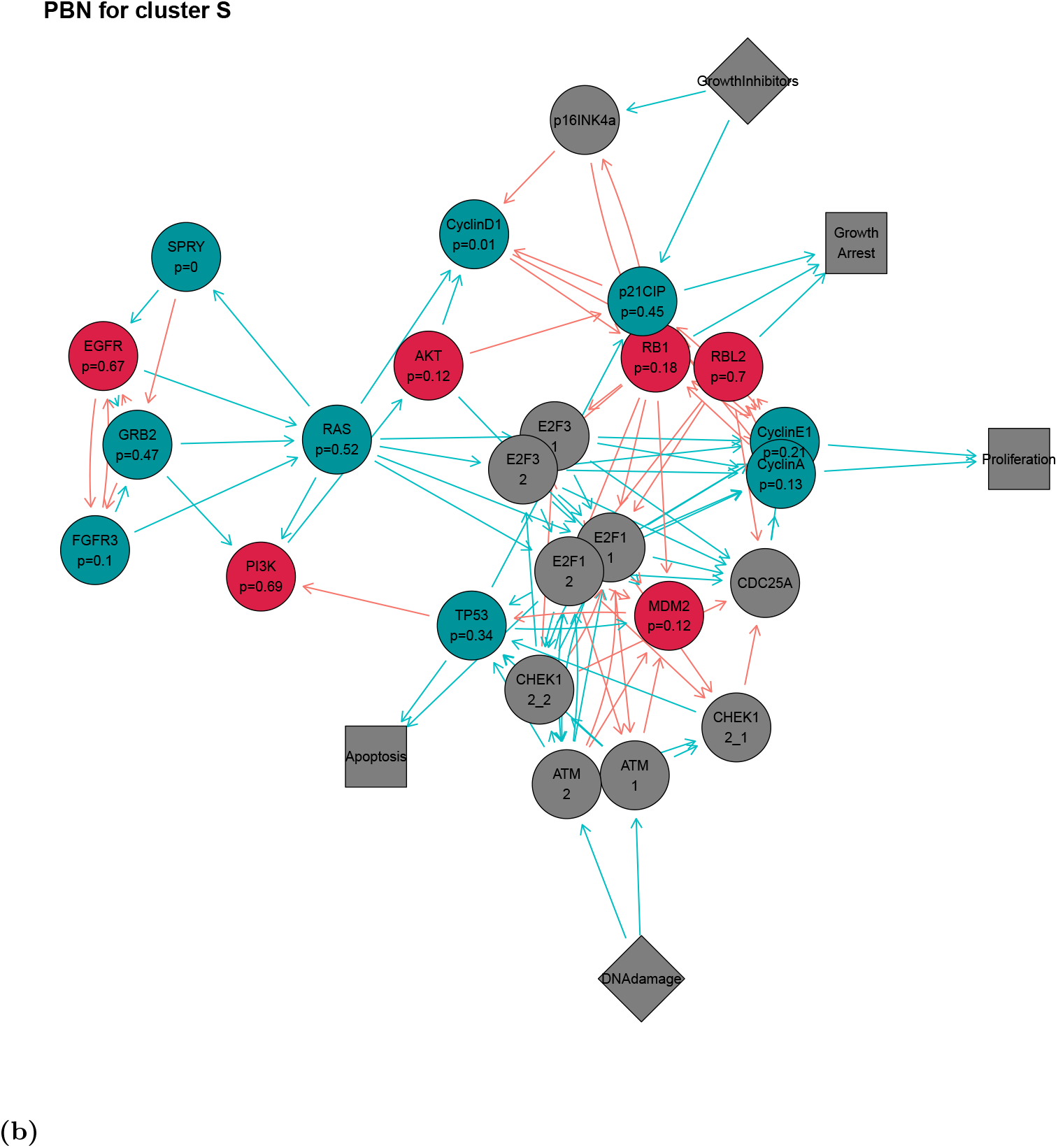
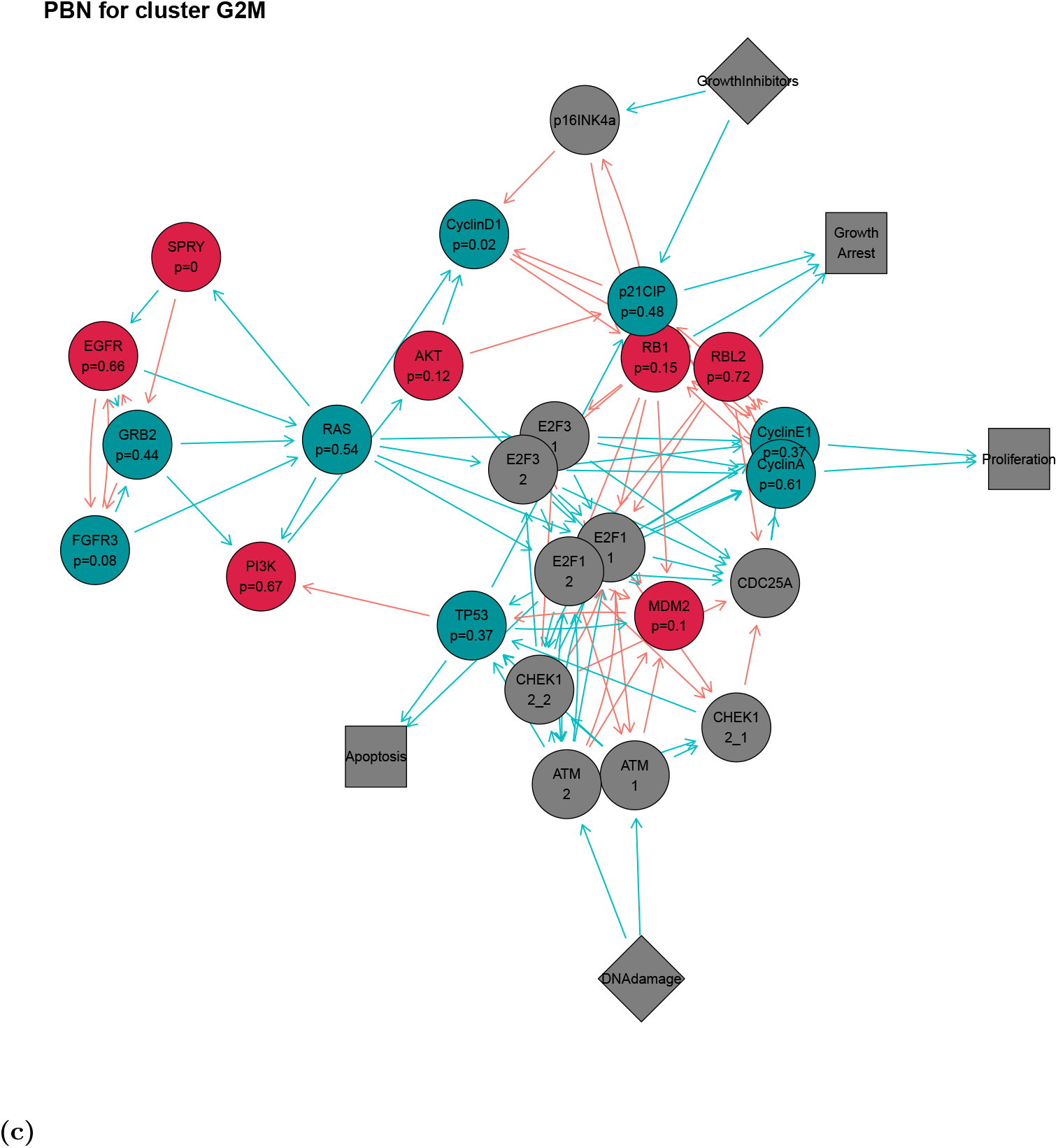
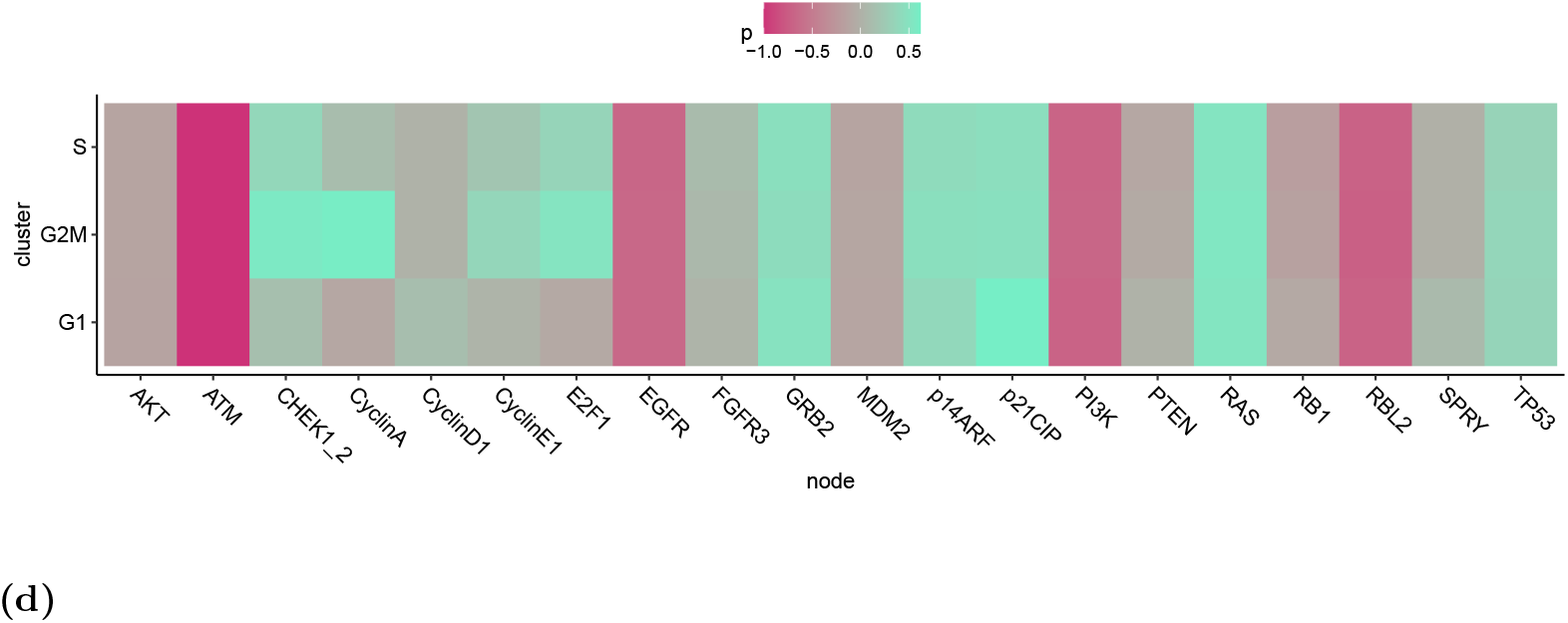
Networks produced by *Ingres* for each of the clusters using the bladder cancer dataset. **a-c**: Each of the PBNs produced by fitting the original Boolean network with the cluster data. **c**: Summary of the networks probabilities as a heatmap. The nodes that are grey in A-C and not present in D weren’t fitted, either because they are an abstraction of several nodes or because they weren’t found by VIPER.

